# Cellular water analysis in T cells reveals a switch from metabolic water gain to water influx

**DOI:** 10.1101/2020.05.11.087767

**Authors:** A Saragovi, T Zilberman, G Yasur, K Turjeman, I Abramovich, M Kuchersky, E Gottlieb, Y Barenholz, M Berger

## Abstract

Cell growth is driven by the acquisition and synthesis of dry biomass and water mass. This study examines the increase of water in T cells biomass during cell growth. We found that T cell growth is initiated by a phase of slow increase of cellular water, followed by a second phase of rapid increase in water content. To study the origin of the water gain, we developed a novel method, Cold Aqua Trap – Isotope Ratio Mass Spectrometry (CAT-IRMS), which allows analysis of intracellular water isotope composition. Applying CAT-IRMS, we discovered that glycolysis-coupled metabolic water accounts on average for 11 femtoliter (fL) out of the 20 fL of water gained per cell during the slow phase. At the end of the rapid phase, before initiation of cell division, a water influx occurs, increasing the water level by three-fold. Thus, activated T cells switch from acquiring metabolic water to incorporating water from the extracellular medium. Our work provides a method to analyze cell water content and an insight into the way cells regulate their water mass.

## Introduction

Cell growth requires cells to accumulate mass and to increase in physical size as a prerequisite step in development and proliferation (Ginzberg *et al*, 2015; Schmoller & Skotheim, 2015). To grow, cells acquire and synthesize additional dry biomass; proteins, lipids, carbohydrates and nucleic acids (Lloyd, 2013), as well as increase their water mass both as free and hydrated molecules (Koivusalo *et al*, 2009; Ho, 2006; Alberti, 2017; Platonova *et al*, 2015). However, how the gain of water mass is achieved, whether primarily due to metabolism or to the change in balance between influx and outflow from the external medium, remains unknown. This has been a challenging problem to study as establishing the source of additional water molecules in cells requires both the differentiation of the intracellular fraction from the background extracellular media and the tracing of the source of water molecules added to the cells. This is important because cellular water quantity affects intracellular chemistry as well as cell intracellular viscosity and mechanics.

Previous attempts to understand the regulation of size, volume and mass in living cells have focused on changes in total cellular mass. This is best assessed using an inertial pico-balance. A recent study, in which single cells were weighed using a pico-balance, revealed that the mass of cells rapidly fluctuates throughout the cell cycle. This study linked these mass fluctuations to the basic cellular processes and water transport (Martínez-Martín *et al*, 2017a). However, although this study described a major leap in our ability to measure dry and water mass changes, it did not enable the direct delineation of cellular water regulation over time. Other efforts to understand cellular water hemostasis were successful in characterizing water uptake rates following osmotic shock (Ho, 2006). Many of these studies used diverse optical methods to establish the dilution rate of the cytoplasm by water influx. Other studies attempted to investigate hydrodynamics at the millisecond level using Raman spectroscopy following acute deuterium exposure (Ibata *et al*, 2011). Likewise, differences in density measurements were also used to extrapolate changes in water mass during cell growth (Delgado *et al*, 2013). However, these indirect methodologies neither trace the source of cellular water mass gains nor directly quantify water influx rates under physiological conditions. Water isotope tracing and NMR were used to investigate organs and whole-body physiological responses in mammals (Pivarnik *et al*, 1984; Schoeller & Van Santen, 1982) and plants (Ionenko *et al*, 2010), but not in the resolution required for analysis at the cellular level. Thus, a method to quantify in cells the correlation between volume changes and water mass gain as well as understanding the origin of water molecules gained during cell growth is lacking.

In this study, we examined the gain of water molecules in growing T cells, as a model for water mass regulation during cell growth. We found that following activation stimuli, T cell water mass increase occurred in two distinct phases, a slow and a fast phase. To establish the contribution of water influx and formation of metabolic water to each of the defined growth phases, we developed a novel method based on the intracellular water isotope composition, called Cold Aqua Trap-Isotope Ratio Mass Spectrometry (CAT-IRMS). We then applied CAT-IRMS to establish the relative contribution of water influx versus metabolic water in each of the defined growth phases. We find that following stimuli, T cells switch between acquisition of metabolic water and increased water influx. During T cell slow growth phase, most water gains result from glycolysis and other metabolic reactions. On the other hand, at the end of the late, fast-growth phase and in proximity to the initiation of cell division, we identified water influx as the predominant source of cellular water gain. We conclude that during activation, T cells switch from metabolic water-based slow-growth to rapid influx-driven water mass increase. The ability to measure both the rate of water gain and the isotope composition of cells’ water mass is an essential tool for the development of specific aquaporin inhibitors as well as for the study of isotope fractionation phenomena in living cells. More importantly, it provides an approach towards a better understanding of the way in which cells determine what size to grow to before dividing.

## Results

### T cell activation is characterized by two distinct cell growth phases

To study water mass incorporation in growing cells, we focused on the well-characterized *in vitro* T cell activation model. Primary naïve T cells are among the smallest mammalian cells, reaching approximately 7 μm in diameter (Tasnim *et al*, 2018). Stimulation of primary T cells leads to the initiation of a tightly coordinated and well-defined cell growth program leading to a significant increase in cell size (Teague *et al*, 1993; Chapman & Chi, 2018). We examined the first 24 hours post stimuli, as during this period T cells go through rapid cell growth independent of cell division (Lea *et al*, 2003).

To explore the dynamics of T cell growth following stimuli, we activated primary T cells *in vitro* using a combination of anti-CD3 and anti-CD28 antibodies. T cell size was approximated using flow cytometry at different time points post stimuli. Analysis of forward scatter signal suggested a two-phase growth dynamic; early slow and late fast growth phases (Figure 1a). Forward scatter signal is a relative parameter and does not always correlate with cell size (Koch L *et al*, 1996). To reach a more robust volume measurement, stimulated T cells were analyzed using a flow cytometer capable of detecting cell electronic volume (EV) based on changes in impendence, in accordance with the Coulter Principle. In line with the forward scatter signal, EV measurements demonstrated a two-phase growth dynamic. In the first slow growth phase, the initial twelve hours following stimuli, average volume per T cell increased from 174.3 to 228.3 fL, ∼4.4 fL/h. During the late fast growth phase, from 12-24 hours post stimuli, average volume per T cell increased from 228.3 to 392.5 fL, at a faster rate of ∼13.96 fL/h (Figure 1b). Primary T cells are a heterogeneous population composed of both CD4+ and CD8+ T cells. We therefore decided to investigate the growth rate of CD8+ and CD4+ T cells separately. Stimulated CD4+ or CD8+ T cells were analyzed using EV flow cytometry. Both T cell types substantially accelerated cell growth at the late phase between 12 and 24 hours post stimuli. Notably, CD8+ T cells, which comprise a third of the total T cell population, accelerated their growth to a larger extent at 12 hours compared to CD4+ T cells, the majority of T cell population. During the late growth phase, the average CD8+ T cell increased by ∼19.8 fL per hour in comparison to ∼12.5 fL/h for the average CD4+ T cell (Figure 1c-d). Together these results demonstrate that at 12 hours following stimuli, T cells switch from a slow to a fast growth program.

**Figure 1:**
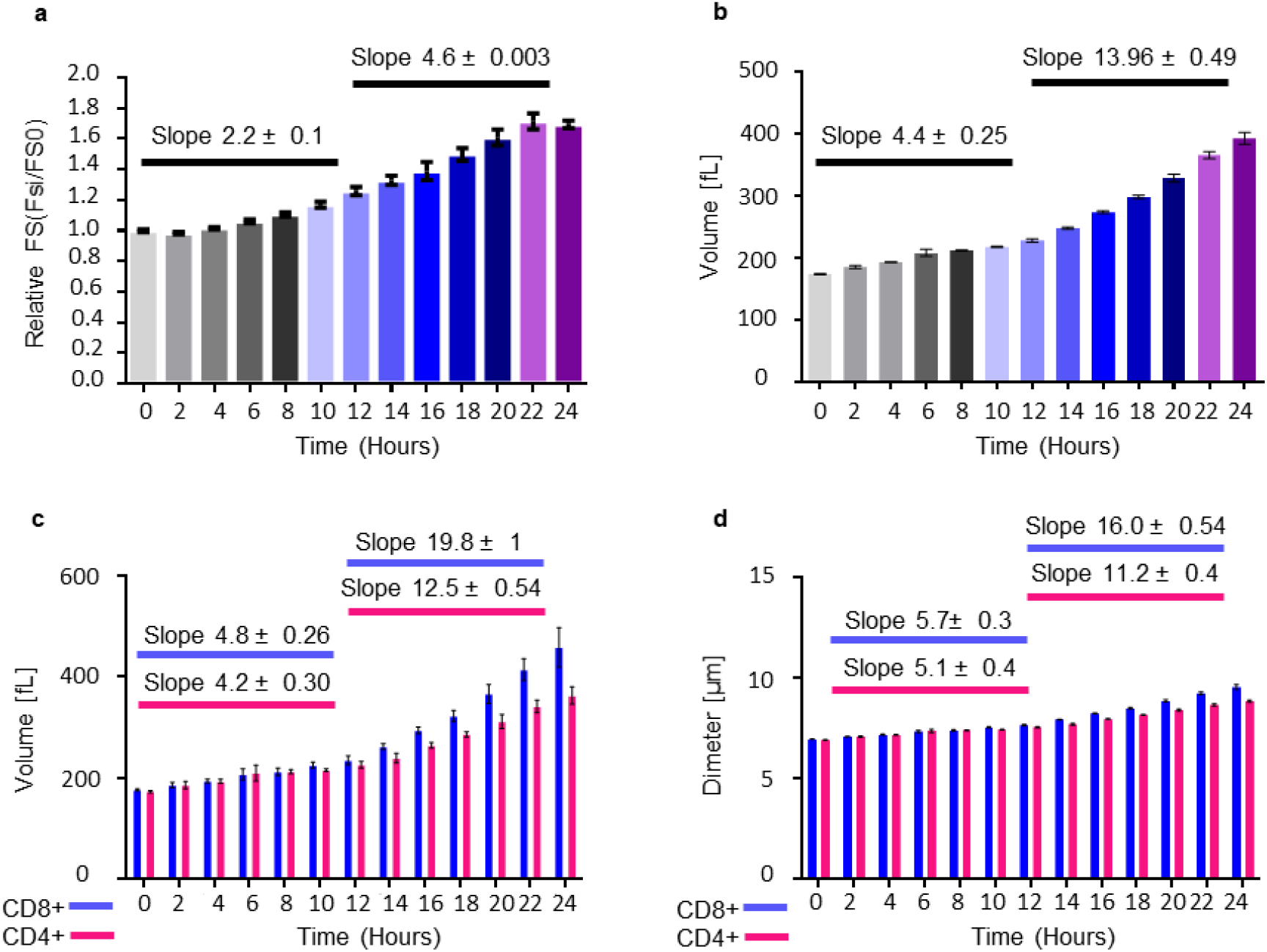
T cell activation is characterized by two distinct cell growth phases: **a-e**. Mouse T cells were activated using anti CD3 and CD28 antibodies and were analyzed by flow cytometer at the indicated time points after stimuli. Bar graphs showing; **a** Relative forward scatter (FS), the ratio between FS of activated to naïve T cells. **b** Electronic volume in fL averaged per cell. **c** Electronic volume in fL averaged per one CD8+ (Blue), or CD4+ (Red) T cell. **d** Diameter in micrometer (μm) averaged per one CD8+ (Blue), or CD4+ (Red) T cell (error bars represent s.e.m.)

### T cell growth is associated with a gradual decrease in water to total volume ratio in the early phase followed by a rapid increase in wet to total volume in the late phase

T cell volume is the sum of both dry and water mass. To directly quantify the differences in T cell water mass during activation, we modified a Karl Fisher titration-based protocol used in liposome content analysis (Cohen *et al*, 2012). We reasoned that we will be able to quantify intracellular water by carefully clearing T cell pellets from surrounding water.

To initially measure the gross water volume in cell samples, T cell pellets were carefully cleared from trace water and resuspended in DMSO (Figure 2a-b). We then used an adjusted Karl Fisher coulometric titration protocol to quantify the water mass in the different samples as well as in DMSO blanks (Figure 2c). To derive the gross water mass, the average DMSO signal (blank) was deducted from each sample (Table S1, equation 1). Finally, to convert the mass to volume, we multiplied the water mass obtained with the water density coefficient, ∼1.0 (Table S1 equation 2).

**Figure 2:**
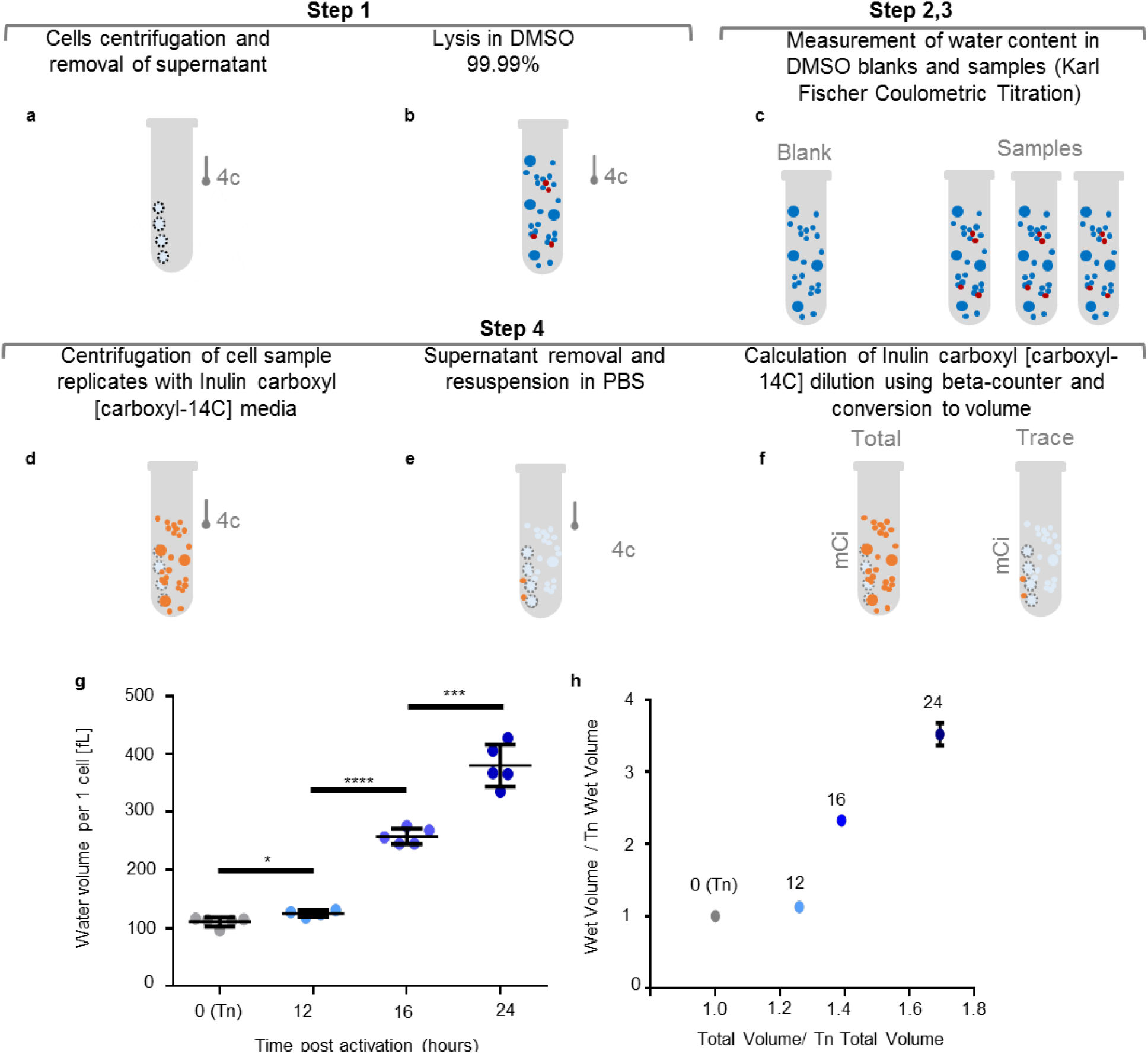
T cell growth is associated with a gradual decrease in wet to total volume ratio in the early phase followed by a rapid increase in wet to total volume in the late phase. **a-c** Schematic of the protocol used for wet volume measurements in activated T cells. **a**. Tubes containing T cell pellets were dried using narrow Whatman papers strips. **b** T cell pellets were dissolved in 99.99% DMSO. **c** The water content in DMSO samples and DMSO blanks were measured using Karl Fischer Coulometric titration. **d-f** Measurement of trace extracellular water in T cell pellet. **d** Cell pellets were resuspended in media containing the cell impermeable Inulin-Carboxyl-^14^C. **e** Following media removal and drying, cells were resuspended in PBS. **f** PBS with trace Inulin-Carboxyl-^14^C was then analyzed in respect to source media using beta counter to derive background volume. **g-h** Mouse T cells were activated using anti CD3 and CD28 antibodies, collected at the indicated time points and analyzed using the Kerl Fisher Coulometric titration protocol. **g** Dot plot presenting averaged, water volume minus trace extracellular water, per one T cell, in fL. **h** correlation between T cell normalized volume and wet volume at different time points (T test significance, * ≤ 0.05, *** ≤ 0.001, **** ≤ 0.0001, error bars represent s.e.m.)

We next attempted to identify the background signal resulting from extracellular water molecules. T cells were suspended in media containing the cell impermeable Inulin-Carboxyl-^14^C (Figure 2d). Cell pellets were then carefully cleared from trace water and resuspended in PBS (Figure 2e). To calculate the extracellular background level, we measured the beta radiation emitted from the PBS used to resuspend the cells (**trace**) with respect to the one from the original Inulin-Carboxyl-^14^C -based media (**source**) (Figures 2f and S1a). To control for other possible background signal sources, we repeated the procedure with empty tubes and also measured the signal from untreated cell samples (Figure S1b). Using these measurements, we could derive the indirect average extracellular water background level per sample by multiplying the dilution factor (mCitrace/mCisource) by the volume of the source sample (Figure S1c and Table S1, equation 3). By combining the background volume quantification with the growth volume measurements and accounting for the number of cells in the sample, we were able to derive the average wet volume per cell in the sample (Table S1, equations 4 and 5).

We then applied this protocol (Figure 2a-f) to quantify the intracellular water mass fraction at several time points during the course of T cell activation. Our measurements indicated that on average, naïve T cells contain ∼111 fL water per cell. Following stimuli, T cell absolute volume increased to 125 fL, 258 fL and 380 fL at 12, 16 and 24 hours, respectively (Figure 2g). Importantly, T cell relative wet volume decreased at the slow growth phase from ∼60% to ∼55% of total volume. In contrast, in the fast phase, the rapid increase in cell wet volume translated to an increase to over 95% water in respect to total cell volume (Figure 2h, S1d). Thus, T cells slow growth phase is associated with a relative decrease in cellular water quantity, while in the fast growth phase, a substantial increase in relative wet volume. These findings suggest that the two well-defined T cell growth phases are underlined by different water mass acquisition mechanisms.

### CAT-IRMS, a direct and robust method for quantifying and identifying the source of cell water mass gain

To identify the source of water gained by T cells during each growth phase, we developed a strategy for intracellular water tracing. Inspired by water tracing studies in plants and bacterial cells (Kreuzer-Martin *et al*, 2005; Walker *et al*, 2001), we designed a three-step protocol for measuring both water influx into cells as well as *de novo* metabolic water. We thought that by a) exposing cells for short periods to labeled media; b) trapping the intracellular fraction by low temperature; and c) analyzing the isotope ratios in the trapped media, we could detect the cellular water isotope composition and derive both the source of and the rate of water gained by the cells (Figure S2a-f). We named this method CAT-IRMS. To trace the entry of extracellular water to the cytoplasm, we chose to treat the cells with water labeled with heavy oxygen isotope (H_2_^18^O)-based medium (Figure S2a). Alternative markers such as deuterium and tritium may diffuse from the labeled solute to various other molecules via proton hopping (de Grotthuss, 2006). Likewise, deuterium-based medium was shown to be toxic to key eukaryotic cellular process including the function of ATP synthase (Olgun, 2007; Nelson & Trager, 2003). In contrast, H_2_^18^O-based labeled medium is inert and remains stable in diverse temperatures (Easteal *et al*, 1984).

A major challenge in isolating the isotope composition of cellular water mass is to separate the intracellular fraction from the extracellular H_2_^18^O-based treatment media. We reasoned that by rapidly cooling cells to a temperature above freezing, we would be able to trap the aqueous content of cells. This is expected because of the thermodynamic effect of low temperature on the rate of osmosis across the cell membrane. In addition, several studies suggest that under temperatures approaching zero Celsius, aquaporins (AQP), the cells water channels (Preston *et al*, 1992), are also partially inhibited (Ionenko *et al*, 2010; Aponte-Santamaría *et al*, 2017). Importantly, because water molecules in eukaryotic cells are locked in a tight sponge-like filament network, the reduction in energy mediated by cooling is expected to cage intracellular water molecules independent of membrane permeability (Sachs & Sivaselvan, 2015). Thus, by cooling the cells, the isotopic composition of intracellular water could then be used to derive both the source and the rate by which cells gained water mass (Figure S2b). To release the trapped intracellular water mass for isotope analysis, a measured volume of double-distilled water (DDW) was added followed by several freeze-thaw cycles and a sonication protocol (Figure S2c). Several control samples were collected and analyzed in parallel to cell samples to account for possible system noise. These included a sample from the labeled media following the treatment (treatment samples) for H_2_^18^O concentration measurement; a sample from the last cold wash media (background) to account for leaks and extracellular contaminants; a sample from the DDW media used to lyse the cells as a baseline. Cell lysate (samples), as well as the specified control samples, were then sent for IRMS analysis (Figures S2 d-f). Since IRMS is fed with gas, a stoichiotransfer was conducted by exposing the samples in close tubes to a gas mixture containing 99.4% helium and 0.6% carbon dioxide for 48 hours at room temperature (Figure S2d). This allows enough time for the oxygen in the carbon dioxide gas to equilibrate with the oxygen in the water. The gas samples containing the oxygen isotope signature was then measured by IRMS (Figure S2e). To assess the minimum sample volume required for stoichiotransfer, we analyzed gas extracts following 48 hours of exposure to decreasing volumes of premeasured H_2_O TW17 isotope standard. Notably, samples greater than 100 µl produced results consistent with the standard, suggesting that volumes ≥ 100 µl are suitable for stoichiotransfer (Figure S3a). IRMS-derived ratios from the different samples were standardized in respect to the Standard Mean Ocean Water (SMOW) isotope ratio (Figure S2f). The measurement of the ^18^O/^16^0 ratio, expressed as d ^18^O for water samples, was made following the CO_2_ equilibration method (Goldsmith *et al*, 2017; Ayalon *et al*, 1998). The results from the standardized IRMS measurements could then be used to derive the total volume of cellular H_2_^18^O in fL and the flux rate, H_2_^18^O*min^-1^ (Table S1, equations 6-9).

To test the robustness and accuracy of CAT-IRMS, we applied our methodology to a number of cell types under diverse conditions. We next examined the influence of H_2_^18^O concentration on the signal level received from cell samples. To assess the limit of detection, Hep2G cells were incubated in media containing 1%, 5% and 50% H_2_^18^O and then analyzed via CAT-IRMS (Figure 3a). Background measurements from the final cell wash were all comparable to the source PBS signal level and 5 permil lower than the sample signal of the cells incubated in 1% H_2_^18^O-enriched media (Figures 3b and S3b). In contrast to background measurements, cell sample measurements produced strong signals, in proportion to the H_2_^18^O concentrations in the incubation media (Figure 3b, S3c). To relate our results to cell number and incubation time, measurements were converted and averaged to H_2_^18^O volume in fL per cell (Figure 3c) and as flux per minute (figure 3d). Under physiological conditions, the accumulation of H_2_^18^O signal in resting cells is anticipated as both growing and quiescent cells are subject to rapid fluctuation in cell density (Martínez-Martín *et al*, 2017b). Due to the concentration of H_2_^18^O, the probability of labeled water to enter the cell is higher than the probability of outflow, so resting cells are expected to accumulate a signal.

**Figure 3:**
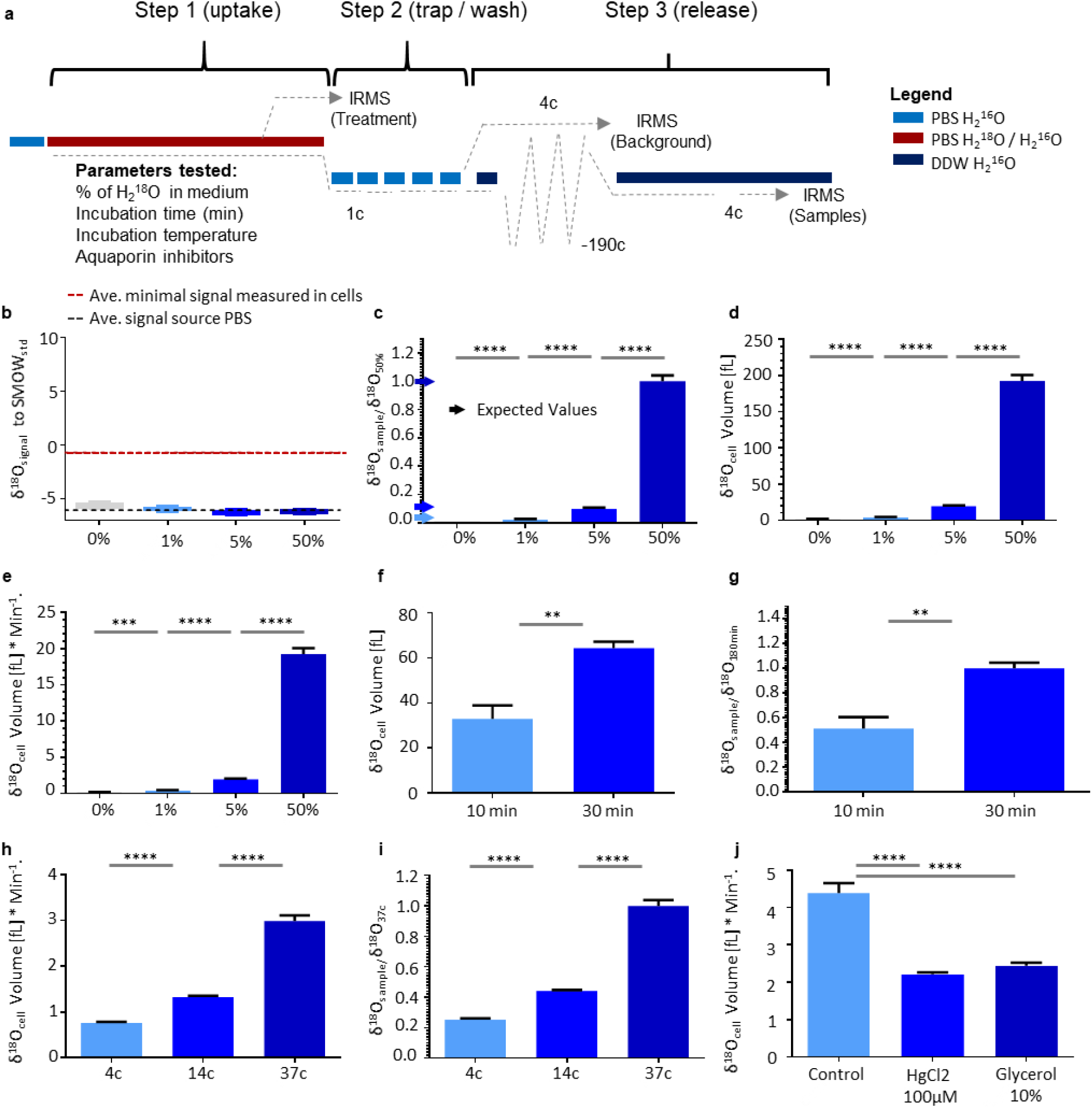
CAT-IRMS, a direct and robust method for quantifying and identifying the source of cell water mass gain. **a** Schematic of overall CAT-IRMS calibration experiments. **b-e**. Hep2G cells were cultured in PBS containing the indicated % of H_2_ ^18^O for 10 minutes. **b** Signal levels in background; d^18^O in the wash media used in the last washing step. Black line represents the average signal in PBS used. Red line represents the average signal in the cells incubated in 1% H ^18^O (showed in c). **c** Relative d^18^O; d^18^O in the sample divided by the average d^18^O in the cells that were incubated in 50% H_2_ ^18^O. Arrows represent the expected values in each concentration group. **d** Averaged H_2_ ^18^O volume in fL per cell. **e** H_2_ ^18^O volume averaged per one cell, per minute (volume of H ^18^O per cell divided by 10 minutes incubation time). **f-g** Hep2G cells were cultured in PBS containing 10% H ^18^O for indicated time intervals. **F** H_2_ ^18^O volume averaged per one cell. **g** H_2_ ^18^O of a sample relative to H_2_ ^18^O in the average sample incubated for 30 minutes. **h-i** Mouse peritoneal macrophages were cultured in PBS containing 50% H ^18^O for 10 minutes at the indicated temperatures. **h** H_2_ ^18^O volume averaged per cell, per minute. **i** H_2_ ^18^O volume in a sample relative to the average H_2_ ^18^O volume in samples that were incubated at 37c. **j** H_2_ ^18^O volume averaged per cell, per minute in peritoneal macrophages incubated in PBS containing 50% H_2_ ^18^O for 10 minutes at 37°C in the absence or presence of the indicated aquaporin’s inhibitors. (T test significance, ns > 0.05, ** ≤ 0.01, *** ≤ 0.001, **** ≤ 0.0001, error bars represent s.e.m.)

To assess the effect of incubation time on the H_2_^18^O signal, Hep2G cells were cultured in media containing 10% H_2_^18^O for 10 or 30 minutes and analyzed for oxygen isotope composition using CAT-IRMS. As expected, cell samples incubated for 30 minutes had a substantially higher H_2_^18^O signal in comparison to cell samples incubated for 10 minutes (Figures 3f-g, S3d). To examine the relevance of CAT-IRMS to primary cells and the effect of temperature on water flux, mouse peritoneal macrophages were cultured in PBS containing 50% H_2_^18^O for 10 minutes at different temperatures. In line with previous reports (Aponte-Santamaría *et al*, 2017; Ionenko *et al*, 2010; Soveral *et al*, 2006), signal from samples of cells incubated in 4°C showed a 5-fold signal reduction in respect to cells cultured in 37°C (Figure 3h-i, S3e). Importantly, the reduction in the signal under low temperature indicates that the potential signal lost during the cold wash cycles step in CAT-IRMS is negligible. AQPs are integral membrane transporters that facilitate water flux across cell membranes (Preston *et al*, 1992). To measure AQP-dependent water flux, mouse peritoneal macrophages were incubated in media containing 50% H_2_^18^O, with or without AQP inhibitors, mercury (II) chloride (Abir-Awan *et al*, 2019; Nicchia *et al*, 2000) or glycerol, to induce subtle hypertonic pressure (Groot & Grubmüller, 2001). As expected, CAT-IRMS analysis of samples treated with AQP inhibitors showed a strong reduction in H_2_^18^O signal in respect to untreated cells (Figure 3j, S3f). Collectively, our measurements indicate that CAT-IRMS is a robust and accurate method, appropriate for analysis and tracing of cellular water flux, synthesis, source and efficiency of AQP inhibitors.

### T cells switch from metabolic water gain during the slow growth phase to water influx at the fast growth phase

The sponge model suggests that in eukaryote cells with dense cytoskeleton, the regulation of intracellular water is not a simple function of extracellular media tonicity (Sachs & Sivaselvan, 2015; Platonova *et al*, 2015). We therefore examined how changes in media tonicity affect naïve, activated T cells volume in respect to Red Blood Cells (RBCs). Incubation of RBCs and resting naïve T cells hypotonic media, led to a significant inflation in cell volume (Figure 4a-b). In contrast, T cells activated in hypotonic media for 12, 16 and 24 hours didn’t demonstrate a significant increase in volume in respect to normal media control (Figure 4c). These findings suggest that cellular water mass growth reflects a tightly regulated mechanism, linked to membrane permeability, the ability of the cytoplasm to absorb water and biogenesis of de novo water molecules by cellular metabolism.

**Figure 4:**
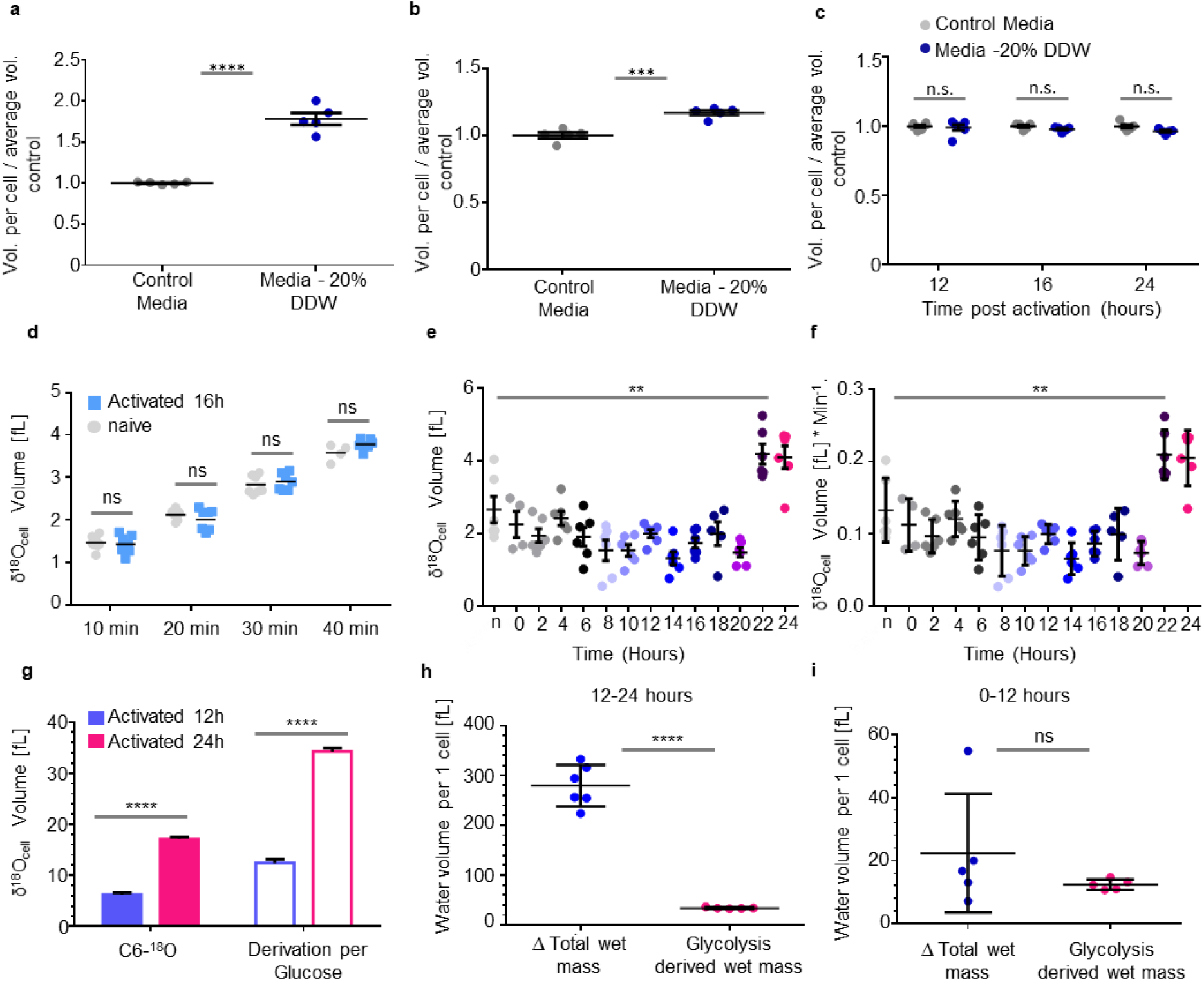
T cells switch from metabolic water gain during the slow growth phase to water influx at the fast growth phase. **a** Human red blood cells were incubated for 12 hours in full media (grey) or media containing 20% DDW (dark blue) and then analyzed via EV flow cytometry **b** Splenocytes were incubated for 12 hours in full media (grey) or media containing 20% DDW (dark blue) and then analyzed via EV flow cytometry (T test significance, ns > 0.05 *, *** ≤ 0.001, **** ≤ 0.0001, error bars represent s.e.m.) **c** Splenocytes were activated using anti CD3 and anti CD28 in full media (grey), media containing 20% DDW (dark blue). Following 12, 16 and 24 hours T cells size was compared via EV low cytometry analysis. (Two way ANOVA test) **d** H_2_ ^18^O volume averaged per cell for mouse naïve or 16 h stimulated T cells that were cultured in a medium containing 50% H_2_ ^18^O for indicated time intervals. **e-f** T cells samples were collected at the indicated time points following stimulation, incubated with 50% H_2_ ^18^O for 10 minutes and analyzed by CAT-IRMS. **e** H_2_ ^18^O volume averaged per cell. **f** H_2_ ^18^O volume averaged per cell per minute **g-i** T cells were stimulated in medium containing C-[6-^18^O]glucose for 12 or 24 hours. **g** H_2_ ^18^O volume measured using CAT-IRMS. Empty frames represent derived values for each glucose molecule (two water molecules per one glucose molecule). Calculated glycolysis derived de novo water biogenesis in respect to total change in water volume **h** during the first 12 hours or **i** 24 hours of T cell activation, averaged per cell. (T test significance, ns > 0.05, ** ≤ 0.01, **** ≤ 0.0001, error bars represent s.e.m.)

To directly test whether stimulated T cells increase the influx of water molecules during growth, we applied CAT-IRMS to measure water entry from extracellular media post stimulation. Naïve and activated T cells, 16 hours post stimuli, were cultured for 10, 20, 30 and 40 minutes in media containing 50% H_2_ ^18^O. Interestingly, IRMS measurements detected no significant differences in cellular H_2_ ^18^O content between the naïve, quiescent and stimulated growing cells (Figure 4d, S4a). To quantify water influx across the first 24 hours of activation, stimulated T cells were treated with media containing 50% H_2_ ^18^O at 13 time points post stimuli. Consistent with our previous results, IRMS measurements detected no significant increase in water uptake by activated cells in the first 22 hours post stimulation. At the late activation period, 22-24 hours post stimulation a two-fold increase in the rate of water uptake was detected, suggesting an additional switch in T cell hydrodynamics around 22 hours post stimuli (Figure 4e-f, S4b).

Following stimulation, T cells rewire their metabolism to support cellular growth. We reasoned that *de novo* water biogenesis may contribute to cellular water mass as a byproduct of these metabolic and anabolic reactions. A hallmark of T cell metabolic alteration following activation is a marked increase in glycolysis (Buck *et al*, 2017). At the ninth step of the glycolysis process, the enzyme enolase catalyzes the conversion of 2 phosphoglycerate to phosphoenolpyruvate (PEP). The hydroxyl groups originated from the carbons in position 6 or 3 in the original glucose molecule are then donated to generate water. To investigate the contribution of glycolysis to intracellular water, we first measured the quantity of glucose consumed by T cells during activation via targeted metabolic analysis. At the first 24 hours post stimuli, we observed more than 40% reduction in media glucose concentration relative to source (Figure S4c). Given the number of cells, the glucose consumed may account for *de novo* biosynthesis of 36 fL of water molecules per cell in total, or 1.5 fL per hour.

To directly measure glycolysis-coupled *de novo* water biogenesis in activated T cells, we used CAT-IRMS to measure the production of H_2_ ^18^O from D-[6 ^18^O]-glucose. Stimulated T cells cultured for 12 and 24 hours produced strong H_2_ ^18^O signal reaching ∼5 fL and ∼15 fL for the average cell, respectively (Figure 4g, S4d). To derive the contribution of *de novo* water biosynthesis to cellular water mass and since D-[6 ^18^O]-glucose labeling accounts for only one out of two water molecules generated, we calculated the final values based on a coefficient of 2 (Figure 4g, empty frames). To compare the water generated by glycolysis to the wet volume gained at each growth phase, we analyzed our measurements relative to the water gained in the two distinct growth phases. Our analysis indicated that glycolysis-derived *de novo* water biogenesis has a marginal contribution to wet volume gains during the fast growth phase (Figure 4h). In contrast, at the slow phase, glycolysis-derived *de novo* water biogenesis accounts for the majority of the increase in water volume gained during the first twelve hours (Figure 4i). Furthermore, since water is a byproduct of mitochondrial respiration as well as protein and RNA synthesis, H_2_ ^18^O measurements vastly underestimate the total intracellular *de novo* water synthesized during T cell activation. Taken together our findings provide compiling evidence that during activation, T cells switch from a slow metabolism-coupled water gain phase to fast, flux-based water mass growth.

## Discussion

Cell growth is a critical step in the development and proliferation of cells. It has not been clear how water molecules are attained during cell growth. The abundance of water in the extracellular fluid may suggest that cells obtain additional water mass by increasing the net influx of water. However, such a mechanism would require cells to continually regulate water fluxes with respect to multiple dry mass components. An intriguing alternative is that cells attain water molecules by *de novo* biosynthesis, as a byproduct of metabolic reactions such as glycolysis, respiration and protein synthesis. Thus, the increase of water mass in growing cells may be coupled to the underlying metabolism of cell growth.

CAT-IRMS, an ^18^O-based method developed in our lab, allows direct measurement of the rate of water influx. Analyzing the isotopic composition of intracellular water is useful not only to trace the source of water flux, but also for identifying *de novo* water generated as a byproduct of metabolism. Furthermore, we have demonstrated CAT-IRMS could be applied to different cell lines and primary cells and be used as a standard to trace intracellular water in multiple applications.

Using CAT-IRMS on activated T cells, we identified three different dominant cellular water gain mechanisms (model provided in Figure S5a). Following activation and during the slow growth phase, T cells acquire water as a byproduct of glycolysis and other metabolic reactions. In contrast, at the late fast growth phase, the increase in water could not be explained by either *de novo* water biogenesis or increased influx. Thus, our observations suggest that during the fast growth phase T cells may increase the overall water volume by decreasing water outflow while keeping influx at the basal level. Finally, as cells approach division, we observed a two-fold increase in the rate of water influx. Since the gain of water in the early slow phase is coupled to the rate of specific reactions, our findings suggest the metabolic divergence may lead to differences in the dry-to-water mass ratio between cells, influencing cell fate decisions. Thus, our work opens a path for a better understanding of cellular growth and for studying the way cells determine what size to grow to before dividing.

## Supplementary Figures

**Figure S1:**
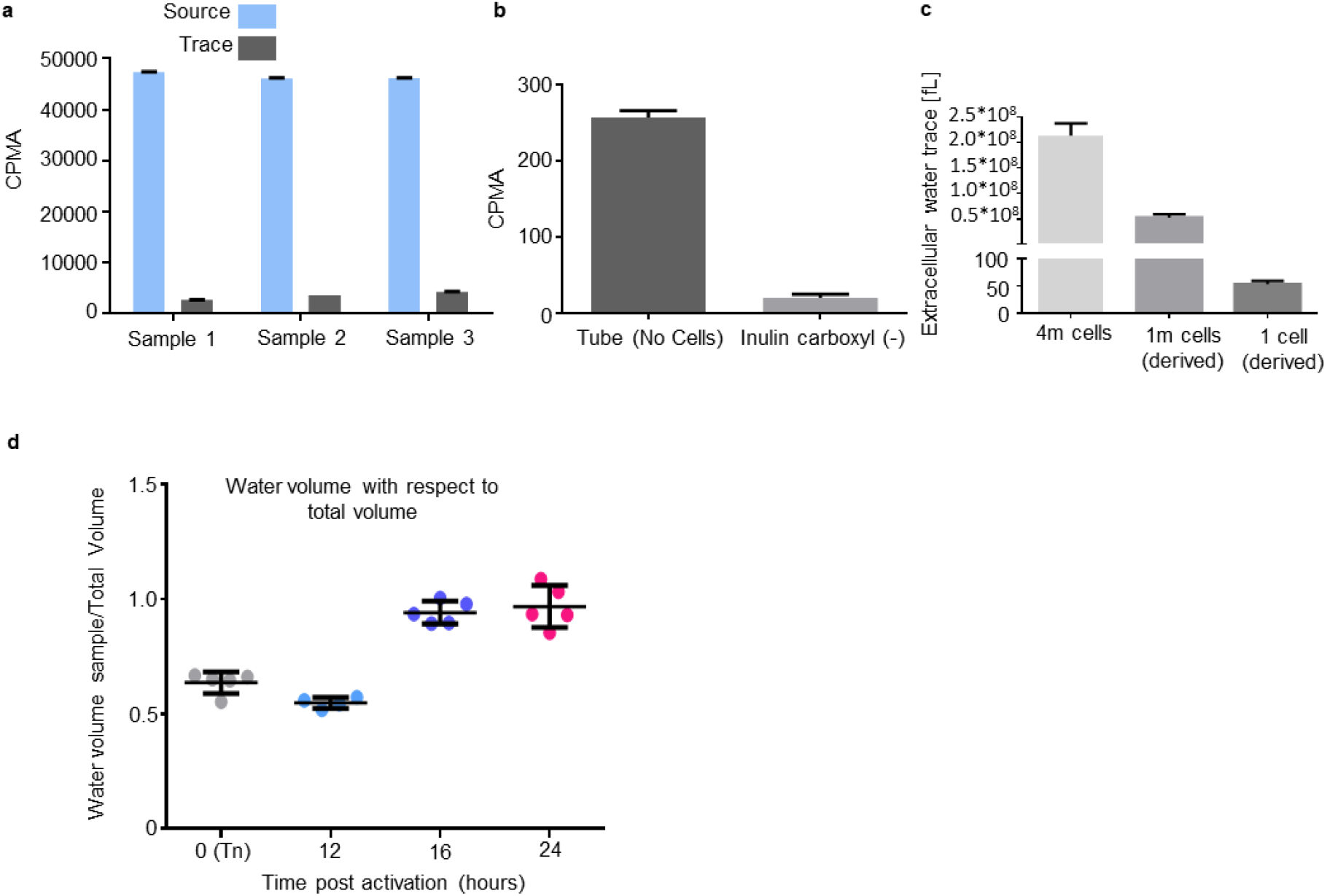
Measurement of trace extracellular water for wet water measurement. **a.** Sample beta counter measurements of source and trace samples (as described in Figure 2f) **b (left)** Source Inulin-Carboxyl-^14^C media was added to empty tubes. Tubes were then dried using thin Whatman strips. Beta counter signal represents the tube background signal (signal independent of cells). **(right)** Beta counter signal of T cell samples in 100ul PBS without Inulin-Carboxyl-^14^C (cell background signal). **c** Trace extracellular background signal calculated for different number of T cells **d**. Water volume in respect to the total volume averaged per one T cell at each time point. (Error bars represent s.e.m.).

**Figure S2:**
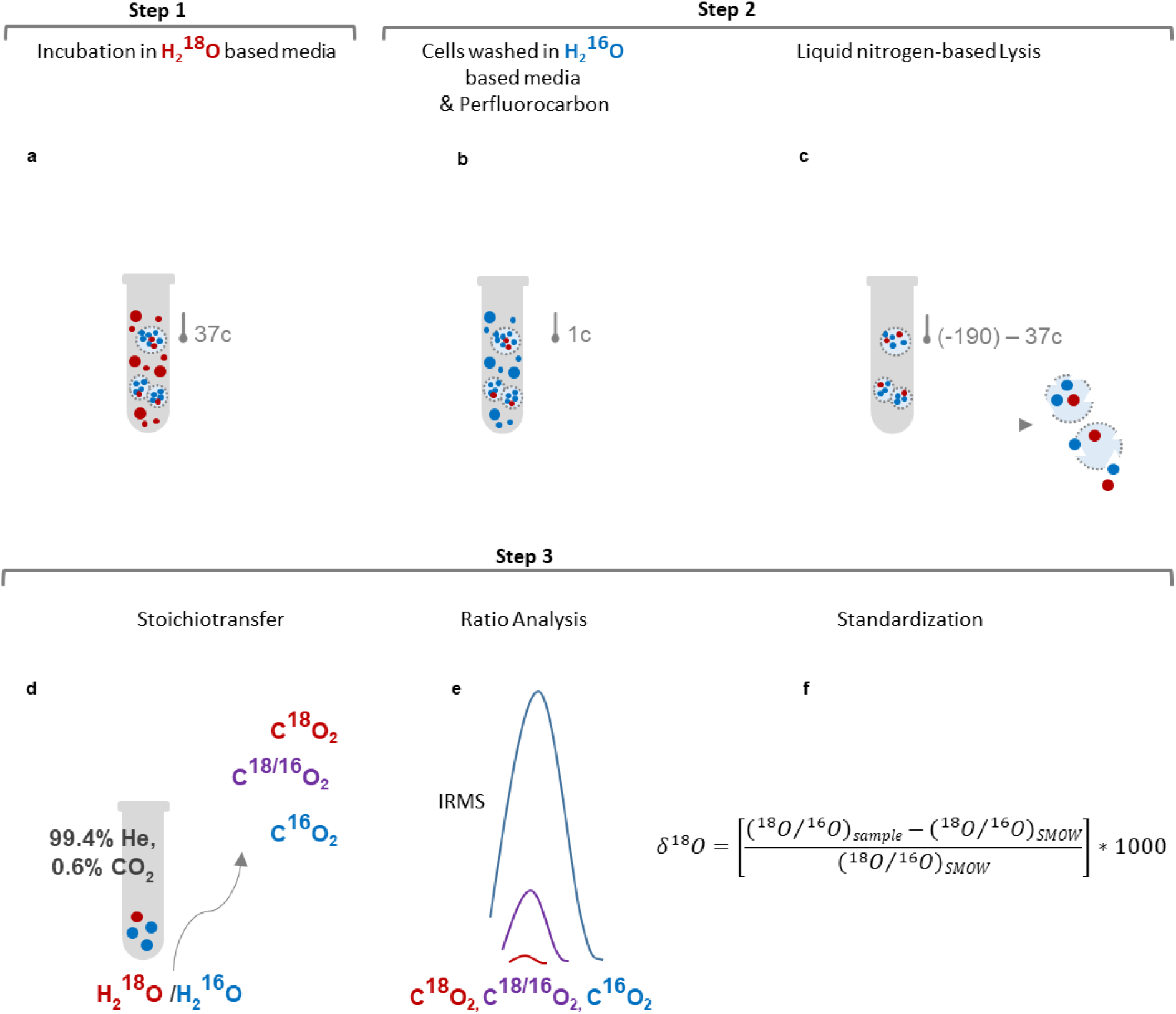
Schematic steps of Cellular Aqua Trap – Isotope Ratio Mass Spectrometry (CAT-IRMS) protocol. Step 1 – **a** Cells are incubated in PBS or culture medium containing different concentrations of H_2_ ^18^O at 37c for an appropriate time intervals. - Step 2 -: **b** Following the incubation, cell samples are rapidly cooled to 1°C and placed on ice in cold room. Cell incubation media is then collected (source). Cells pellets are next washed 5 times with ice cold normal PBS. **c** To release the intracellular water trapped within the cells, 400 µl DDW are then added to the cell pellets followed by 3 cycles of freeze/thaw using liquid nitrogen and sonication. Cell lysates are then collected and sent for IRMS analysis for ^18^O/^16^O ratio measurements. Media from steps 1 (source) and 2 (background) are also collected and sent to IRMS **-** Step 3:-**d** For IRMS analysis, samples were injected into sealed glass tubes filled with 99.4% He, 0.6% CO_2_ for 48 hours for stoichiometric transfer of ^18^O or ^16^O to CO_2_ **e** Gas mixture from each tube is then loaded into the IRMS for measurement **f** ^18^O volume (δ^18^O) is extrapolated by comparing to international H_2_ ^18^O abundance in ocean water using the presented formula (f). *SMOW*: Standard Mean Ocean Water.

**Figure S3:**
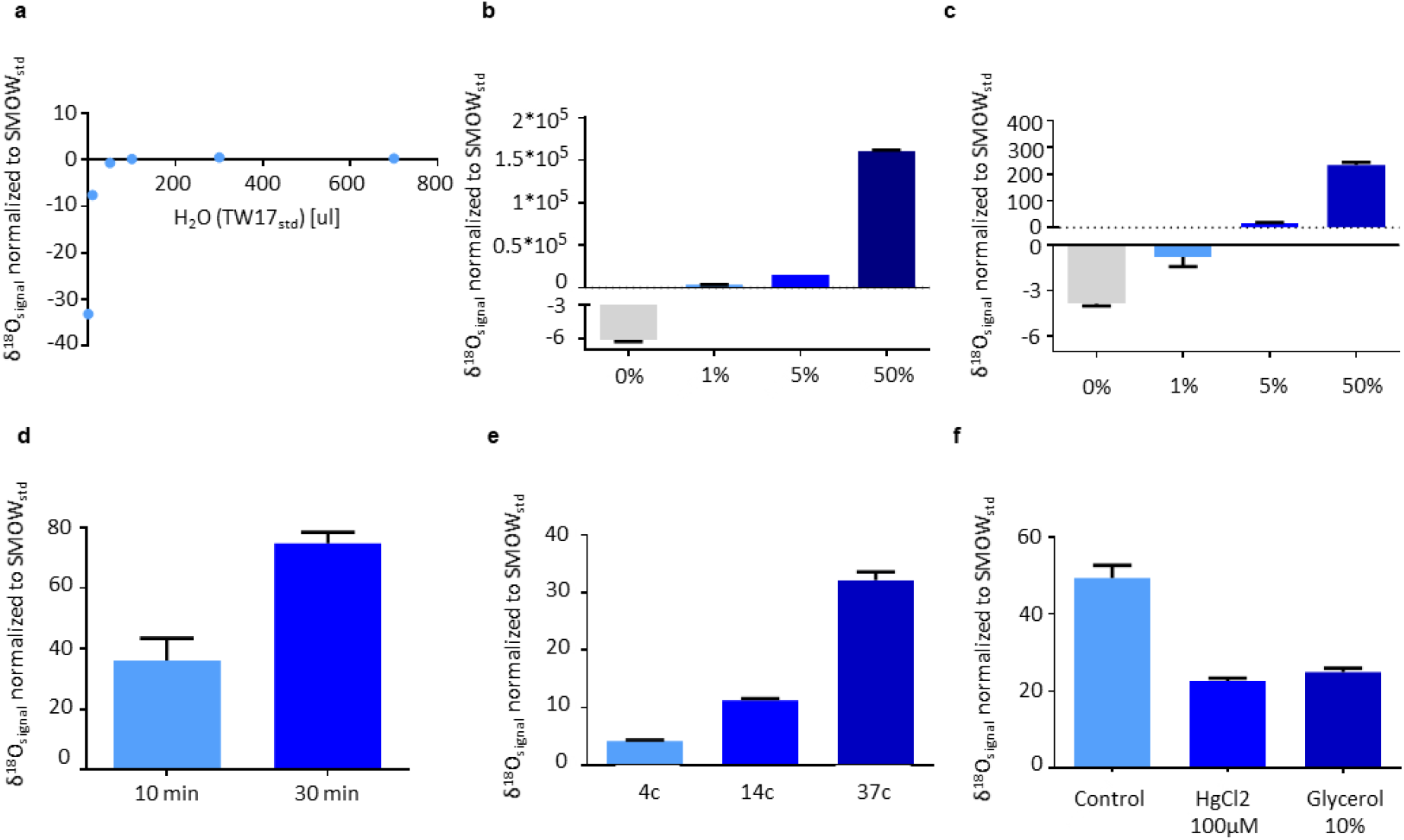
Calibration of CAT-IRMS. **a** Measurement of minimum volume required for stoichiometric transfer in δ^18^O signal normalized to SMOW b-c. Hep2G cells were cultured in PBS containing the indicated % of H_2_ ^18^O for 10 minutes. **b** δ^18^O in the source media used to incubate the cells. Bars represent δ^18^O signal normalized to SMOW. **c** Cell samples δ^18^O signal normalized to SMOW. **d** Hep2G cells were cultured in PBS containing 10% H_2_ ^18^O for indicated time intervals. Bars represent δ^18^O signal normalized to SMOW **e** Mouse peritoneal macrophages were cultured in PBS containing 50% H_2_ ^18^O for 10 minutes at the indicated temperatures. Bars represent δ^18^O signal normalized to SMOW **f** δ^18^O signal normalized to SMOW in peritoneal macrophages incubated in PBS containing 50% H_2_ ^18^O for 10 minutes at 37°C in the absence or presence of the indicated aquaporin inhibitors. (Error bars represent s.e.m.)

**Figure S4:**
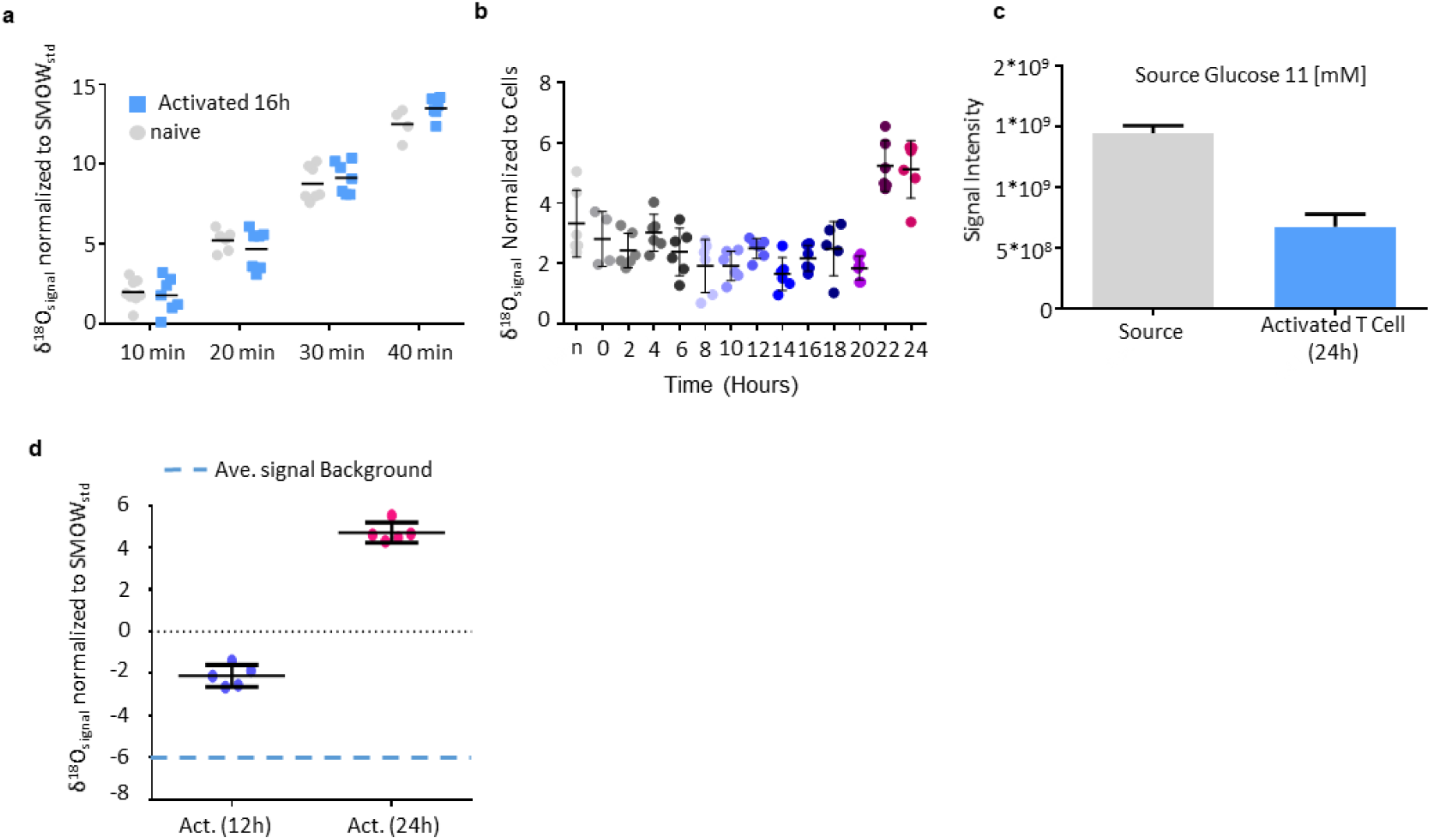
T cells growth is propagated by three distinct water mass gain states. **a** δ^18^O signal normalized to SMOW in samples from mouse naïve or 16 h stimulated T cells that were cultured in a medium containing 50% H_2_ ^18^O for indicated time intervals. **b** T cell samples were collected at the indicated time points following stimulation, incubated with 50% H_2_ ^18^O for 10 minutes and analyzed by CAT-IRMS. Dots represent δ^18^O signal normalized to SMOW **c** Glucose levels as signal intensity in growing media of activated T cells for 24 hours relative to glucose levels in the source media. Glucose levels were measured using LC-MS-MS. **d** Naïve or stimulated T cells were cultured in medium containing D-[6-^18^O]glucose for 12 or 24 hours. Samples were then measured using CAT-IRMS. Dots represent δ^18^O signal normalized to SMOW. (Error bars represent s.e.m.)

**Figure S5:**
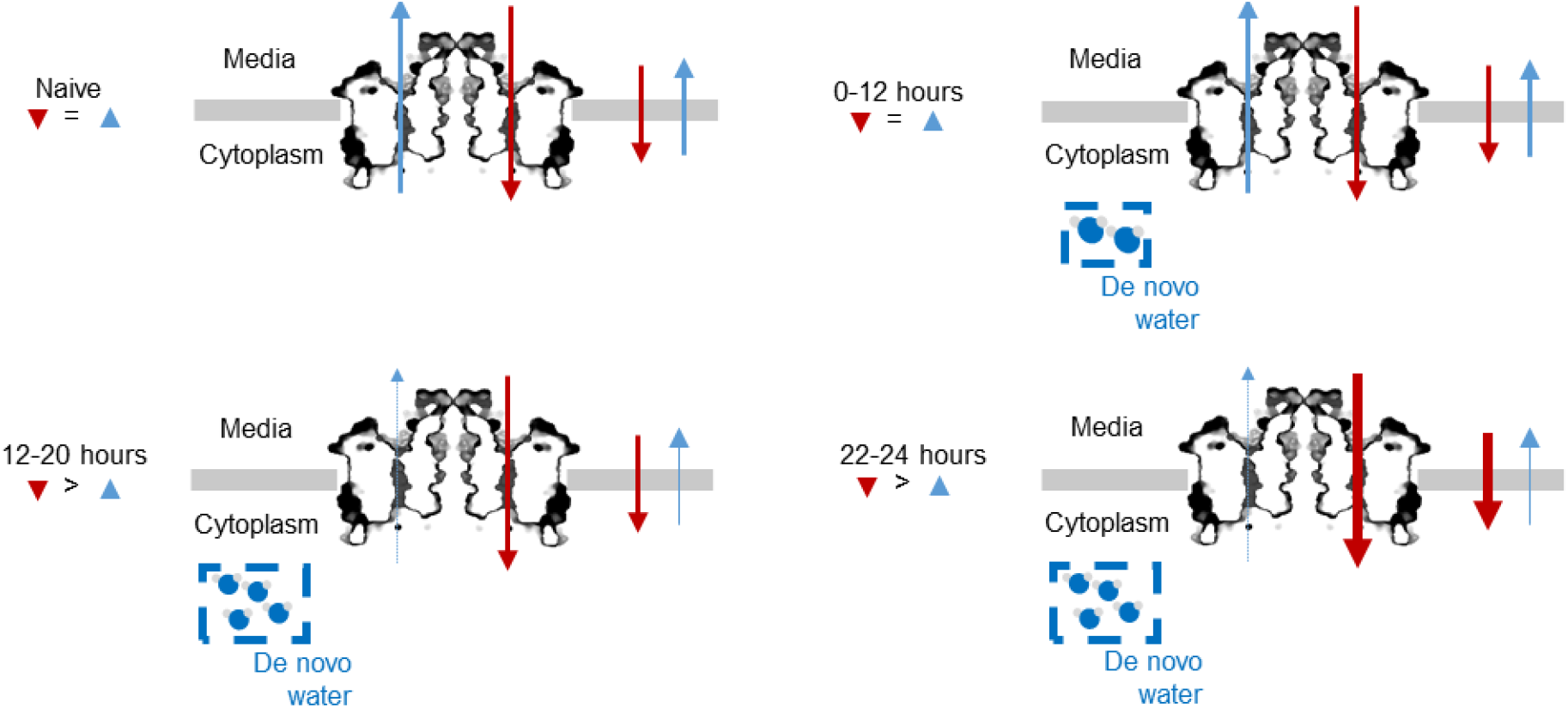
T cells growth is propagated by three distinct water mass gain states. A model representing the different states of cellular water mass gain during T cell activation

**Supplementary Table 1.**

| # | Equation used to | Equation |
| --- | --- | --- |
| 1 | derive the growth sample wet mass | $WetMass_{gross} = KF_{sample} - KF_{blank}$ |
| 2 | convert sample wet mass to wet volume | $WetVolume_{gross} = KWetMass_{gross} * Water_{density}$ |
| 3 | calculate extracellular trace water volume per sample | $WetVolume_{extracellular} = \frac{mCi_{trace}}{mCi_{source}} * Volume_{source}$ |
| 4 | the intracellular wet volume per sample | $WetVolume_{intracellular} = WetVolume_{gross} - WetVolume_{extracellular}$ |
| 5 | average intracellular water volume per cell | $WetVolume_{cell} = \frac{WetVolume_{intracellular}}{Cell_{number}}$ |
| 6 | normalize $\delta^{18}O$ signal to SMOW | $\delta^{18}O = \frac{{}^{18}O/{}^{16}O_{sample} - {}^{18}O/{}^{16}O_{std}}{{}^{18}O/{}^{16}O_{std}} * 1000$ |
| 7 | calculate total sample $H_2^{18}O$ volume | $H_2^{18}O_{volume} = \frac{\delta^{18}O_{sample} - \delta^{18}O_{background}}{500,000} * Sample_{volume}$ |
| 8 | derive $H_2^{18}O$ volume per cell | $H_2^{18}O_{cell} = \frac{H_2^{18}O_{volume}}{Cell_{number}}$ |
| 9 | calculate $H_2^{18}O$ influx per minute | $H_2^{18}O_{flux} = \frac{H_2^{18}O_{cell}}{t_{incubation}}$ |

## Materials and Methods

### Mice

The C57BL/6J (wild-type) mice were from The Jackson Laboratory. Mice were maintained and bred under specific pathogen-free conditions in the Hebrew University animal facilities according to Institutional Animal Care and Use Committee regulations. All mice were maintained on the C57BL/6J background and used for experiments at 8–12 weeks of age.

### Human Samples

Human blood samples were obtained via Shaare Zedek Medical Center Jerusalem, Helsinki committee approval number: 143/14.

### T cells Isolation

CD8+ or total T cells were isolated from spleens with an EasySep(tm) T Cell Isolation Kit according to the manufacturer’s instructions (STEMCELL Technologies).

### Peritoneal macrophage culture

Residential peritoneal macrophages were directly collected from anaesthetized mice by lavage with 5 ml PBS using a 5 ml syringe with 19G needle. When elicited macrophages were needed, the mice were injected intraperitoneally with 1 ml 4% thioglycollate at least 36 hours in advance. Then, 5 × 105 erythrocyte-depleted lavage-derived cells were plated on a tissue culture treated polystyrene 24-well plates (Corning, NY).

### In vitro T cell proliferation assay

Primary isolated T cells or CD8+ were activated in 96-flat-well plates (1 × 10^6^ cells per well) coated with anti-CD3ε (6 µg/ml) and anti-CD28 (6 µg/ml).

### Flow Cytometry and EV

Cells were stained with various conjugated mAbs against cell-surface markers in FACS buffer (PBS containing 1% FBS) for 30 min at 4°C. Stained cells were assayed by Gallios flow cytometer with Kaluza software (Beckman Coulter, Brea, CA) or iCyt Eclipse (Sony) and analyzed by FACS Express 6 (De Novo Software).

### Karl Fisher Titration

Isolated naïve T cells were centrifuged in 1.7 ml tubes. Following careful removal of all visual liquids, pellets were resuspended in DMSO (Sigma 99.99%). Samples were measured in 831 KF Coulometer (Metrohm). For calculation of background noise blank DMSO measurements were performed. To calculate the noise from water leftovers, 10 samples were suspended in PBS containing Inulin-Carboxyl-^14^C. Samples were then centrifuged, and PBS removed. Cells were resuspended in 1ml PBS and Inulin-Carboxyl-^14^C radioactivity was measured to evaluate dilution in respect to standard.

### Targeted metabolic analysis

CD8+ T cells were cultured in anti-CD3/CD28 coated 96-well plate (1 million cells /well), suspended in RPMI supplemented with 10% dialyzed fetal bovine serum and 100 μM alanine. Following 24 hours activated cells were then extracted for metabolomics LC-MS analysis.

Medium extracts: 20 μl culture medium were added to 980 μl of a cold extraction solution (–20°C) composed of methanol, acetonitrile, and water (5:3:2). Medium extracts were centrifuged (10 min at 16,000 g) to remove insoluble material, and the supernatant was collected for LC-MS analysis. Metabolomics data was normalized to protein concentrations using a modified Lowry protein assay.

LC-MS metabolomics analysis was performed as described previously (MacKay *et al*., 2015). Briefly, Thermo Ultimate 3000 high-performance liquid chromatography (HPLC) system coupled to Q-Exactive Orbitrap Mass Spectrometer (Thermo Fisher Scientific) was used with a resolution of 35,000 at 200 mass/charge ratio (m/z), electrospray ionization, and polarity switching mode to enable both positive and negative ions across a mass range of 67 to 1000 m/z. HPLC setup consisted ZIC-pHILIC column (SeQuant; 150 mm x 2.1 mm, 5 μm; Merck), with a ZIC-pHILIC guard column (SeQuant; 20 mm x 2.1 mm). 5 μl of biological extracts were injected and the compounds were separated with mobile phase gradient of 15 min, starting at 20% aqueous (20 mM ammonium carbonate adjusted to pH.2 with 0.1% of 25% ammonium hydroxide) and 80% organic (acetonitrile) and terminated with 20% acetonitrile. Flow rate and column temperature were maintained at 0.2 ml/min and 45°C, respectively, for a total run time of 27 min. All metabolites were detected using mass accuracy below 5 ppm. Thermo Xcalibur was used for data acquisition. TraceFinder 4.1 was used for analysis. Peak areas of metabolites were determined by using the exact mass of the singly charged ions. The retention time of metabolites was predetermined on the pHILIC column by analyzing an in-house mass spectrometry metabolite library that was built by running commercially available standards.

### Isotope Ratio Mass Spectrometry (IRMS)

For the measurement of d^18^O water, clean vacuum vessels were flushed with a gas mixture of Helium (99.6%) and CO2 (0.4%) for 10 min to remove the original atmosphere. After flushing, 0.7 cm^3^ of the sampled water was injected to the vessels and left to equilibrate with the CO2 gas for 48 hours at 25 °C. Oxygen isotopes were measured using a using a Finnigan Gas Bench II extraction system attached to a ThermoFinnigan Delta PLUS XP continuous flow mass-spectrometer. All oxygen isotopic measurements were done in duplicate and reported relative to VSMOW. Four well calibrated internal laboratory standards were used for calibration and a standard was measured every 8 water samples.

### Statistical analysis

The statistical significance of differences was determined by the two-tailed Student’s t-test. Differences with a P value of less than 0.05 were considered statistically significant. Graph-prism software was used for analysis.

## Author contributions

A.S. and M.B. designed and performed research, analyzed data and wrote the manuscript; T.Z., G.Y., K.T., I., MK., E.G. and Y.B. performed research.

## Acknowledgments

This work was supported by grants from the Israel Science Foundation grant No. 1596/17 and German-Israeli Foundation grant No. I-224-414.11-2017.

